# Morphometry and mechanical instability at the onset of epithelial bladder cancer

**DOI:** 10.1101/2023.08.17.553533

**Authors:** Franziska L. Lampart, Roman Vetter, Yifan Wang, Kevin A. Yamauchi, Nico Strohmeyer, Florian Meer, Marie-Didiée Hussherr, Gieri Camenisch, Hans-Helge Seifert, Cyrill A. Rentsch, Clémentine Le Magnen, Daniel J. Müller, Lukas Bubendorf, Dagmar Iber

## Abstract

Malignancies of epithelial tissues, called carcinomas, account for the majority of cancer cases. Much cancer research has focused on genetic alterations and their relation to different carcinoma phenotypes. Besides a rewiring in the signalling networks, carcinoma progression is accompanied by mechanical changes in the epithelial cells and the extracellular matrix. Here, we reveal intricate morphologies in the basement membrane at the onset of bladder cancer, and propose that they emerge from a mechanical buckling instability upon epithelial overgrowth. Using a combination of microscopy imaging of the mouse and human bladder tissue, elasticity theory, and numerical simulations of differential growth in the bladder mucosa, we find that aberrant tissue morphologies can emerge through stiffness changes in the different mucosa layers. The resulting thickening, wrinkles and folds exhibit qualitative and quantitative similarity with imaged early papillary tumors and carcinomas *in situ*. Atomic force microscopy indeed reveals local stiffness changes in the pathological basement membrane. Our findings suggest a mechanical origin of the different carcinoma subtypes in the bladder, which have vastly different clinical prognosis. They might provide the basis for a new line of attack in medical carcinoma treatment and prophylaxis.

## Introduction

Solid tumors that originate from epithelia, so-called carcinomas, are by far the most frequent cancers [1]. Epithelia are thin tissues that line the internal and external surfaces of a body. Watertight tight junctions on their apical side allow them to seal surfaces, while basal adhesion to the stiff basement membrane (BM), a thin layer of extracellular matrix (ECM), provides mechanical stability [2, 3]. A hallmark of tumorigenesis is the local disintegration of the epithelial tissue architecture, which allows cells to breach the BM and metastasise [4]. Deregulated epithelial growth can be directed either inward (endophytic) or outward (exophytic). The direction of epithelial expansion can influence the aggressiveness of tumors such as in bladder, kidney, skin and uterine cervical cancers [5–9], and is taken into consideration when deciding on treatment strategies and post-treatment surveillance [10–13]. How genomic mutations translate into different tumor growth patterns is still largely unknown. Defining the underlying physical drivers of tumor morphogenesis may help to advance diagnostic and treatment approaches.

Bladder cancer (BC), one of the most expensive cancers to manage [14], offers a particularly suitable model system to study the emergence of distinct epithelial morphologies at the onset of carcinogenesis. The bladder is a particularly accessible organ, enabling easy collection and imaging of tissue samples at all stages of cancer development. The risk of urothelial BC invasion into deeper tissue layers depends on the growth pattern [15]. The so-called papillary tumors form fingerlike protrusions either into the bladder lumen (exophytic growth) or, as inverted urothelial tumors, into the subepitheliale connective tissue (endophytic growth). Low-grade papillary tumors have a low risk of progression. Flat localised carcinoma *in situ* (CIS) (planophytic growth), on the other hand, have a high risk for progression to muscle-invasive BC [16, 17]. Moreover, BC stands as a good example of the gap between genotype and phenotype. It is still unclear how mutations translate into papillary or CIS tumor morphologies, despite the notable correlation between growth patterns and specific mutations in these carcinomas [5]. With the N-Butyl-N-(4-hydroxybutyl) nitrosamine (BBN) mouse model [18], the early onset and later progression of BC can directly be observed and investigated. The ability to induce tumors of different morphologies and observe early progression makes BC the ideal system to study the determinants of endophytic vs. exophytic growth.

Tissue curvature, as proposed recently for tubular epithelia in the pancreas of mice [6], can be ruled out as a reliable predictor for the growth of papillary vs. CIS in BC, as the bladder in mice and larger animals is too big, such that the tissue curvature is too low. Another important aspect is tissue mechanics. Tubular mucosa mechanically wrinkles under volumetric growth [19, 20]. The stiffness of the BM has been related to the emergence of either buds or folds in flat skin carcinomas [7]. Tumor-associated cells actively alter the structure and mechanical properties of the ECM [21–23]. Modifications to the tumor ECM impact cancer progression and treatment response [24, 25]. To become invasive and eventually metastasize, carcinomas must breach the BM. While proteases that remodel and degrade ECM components have been known to facilitate BM invasion [22, 26], recent research indicates that mechanical forces can also promote invasion independently of protease activity [27–29]. Despite the known changes in mechanical properties of the ECM [30–33] and ample evidence for the impact of ECM mechanics on cancer progression [22, 23, 30, 32, 34–36], the effect on the tumor morphology is not well understood.

In this article, we apply concepts from continuum mechanics along with three-dimensional (3D) microscopy and stiffness assessments using atomic force microscopy (AFM). We uncover BM morphologies during the early stages of BC that we demonstrate can arise from a mechanical buckling instability, as a result of overgrowth of the urothelial layer and changes in the tissue stiffness. We characterize the morphological hallmarks of different BC subtypes in 3D imaging of human biopsies and mouse tissues and find that their onset can be recapitulated by continuum simulations of an elasto-plastic three-layered tissue undergoing differential growth. Changes in the composition and stiffness of the ECM in BC have already previously been investigated in the context of progression to muscle invasive BC and metastatic BC [37]. We now identify the relative stiffness of the three mucosa layers—the urothelium, the BM, and the lamina propria (LP)—as the main mechanical discriminant between exophytic and endophytic growth upon malignant proliferation of the epithelial cells. Consistent with this theory, AFM on mouse BMs confirms localised softening, which our model predicts to affect the malignant potential of urothelial proliferation.

## Results

### Early onset of bladder cancer visible in the structure of the basement membrane

In order to image the onset of BC, we chemically induced it by subjecting 10-week-old male mice, that express a green fluorescent membrane marker (mEGFP) in the urothelial layer and a red fluorescent membrane marker (m-tdTomato) throughout the rest of the bladder tissue [38], to BBN in the drinking water (Fig. 1A,B, Methods). All bladders from the BBN treatment cohort showed alterations on the inside of the bladder wall 11 weeks after BBN treatment. On a macroscopic level, these alterations appear as nodular structures (Fig. 1B arrow). We encountered no such structures in bladders from the control cohort (Fig. 1C).

**Figure 1:**
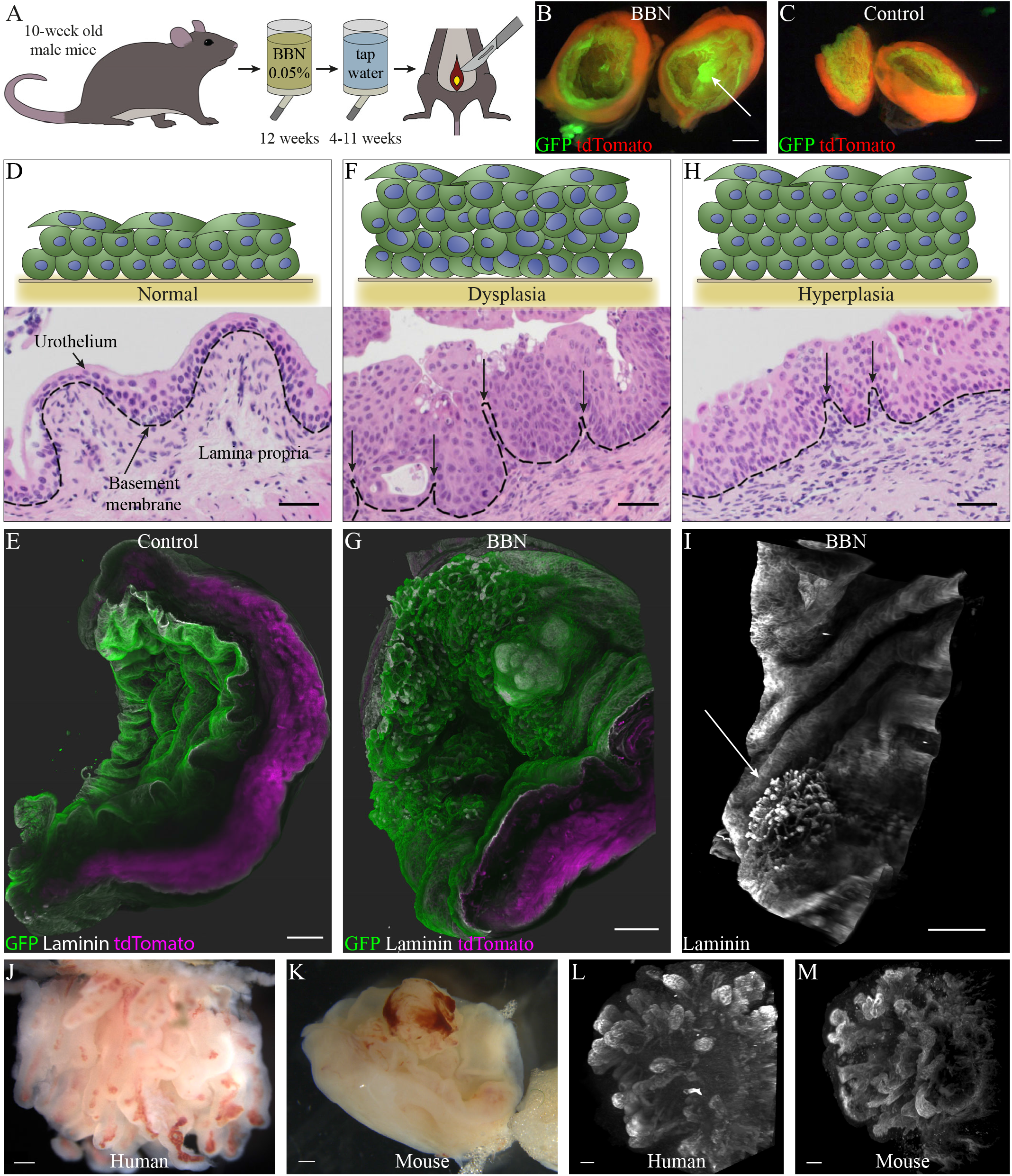
BBN induces changes in the mucosa of mice. (A) BBN treatment regime, (B–C) Macroscopic images of mouse bladders from treatment (B) and control (C) cohort, scale bar: 1 mm, (D) Illustration of a normal urothelium and histopathological section of mouse bladder tissue from the control cohort, scale bar: 50 μm, (E) 3D SPIM images showing the normal folding of a healthy mouse bladder mucosa, scale bar: 500 μm, (F) Illustration of a urothelium with dysplasia and histopathological section of mouse bladder tissue with dysplasia from the treatment cohort week 11 post BBN, scale bar: 50 μm, (G) 3D SPIM images showing the folding patterns of a mouse bladder mucosa from the treatment cohort at week 4 post BBN, scale bar: 500 μm, (H) Illustration of a urothelium with hyperplasia and histopathological section of mouse bladder tissue with hyperplasia from the treatment cohort week 4 post BBN, scale bar: 50 μm, (I) BM of a mouse tissue from the treated cohort at week 4 post BBN, scale bar: 500 μm, (J–K) Biopsy of a human papillary tumor (J) and a BBN induced tumor in mice 11 weeks post BBN (K), scale bar: 500 μm, (L–M) BM structure in a human pTa tumor (L) and mouse bladder tumor (M), scale bar: 100 μm

The urothelium in a healthy mouse bladder is about three layers thick, and in the void bladder, the mucosa, consisting of the urothelium, the BM and LP, folds in a regular pattern as seen in histopathological sections and the 3D reconstructions of single plane illumination microscopy (SPIM) z-stacks (Fig. 1D and E). At 11 weeks after BBN treatment, various types of urothelial neoplasms can be detected in the histopathological sections, such as dysplasia (Fig. 1F), papillary tumors and CIS (Fig. S1A and B), with a thickening of the urothelium and abnormal-looking cells. In the dysplastic tissue, narrow folds of the BM and LP are visible (Fig. 1F, arrows), which are much smaller in diameter than would be anticipated from the mucosa’s typical bulging in a void bladder (Fig. 1D and E). These changes in the mucosa are also visible in 3D reconstructions of SPIM z-stacks from BBN cohort mice with neoplastic growth (Fig. 1G). Similar, but smaller, structures can be seen in bladders with hyperplasia as early as 4 weeks after treatment (Fig. 1H, arrows). Here, the urothelium surrounding these structures shows hyperplastic growth with a thickened urothelium, but otherwise has normal cytological appearance (Fig. 1H), demonstrating that these narrow folds already arise in precancerous stages. The morphological changes of the mucosa become especially apparent in the BM, where a localized buckling pattern emerges that is noticeably different from the otherwise rather smooth BM (Fig. 1I, arrow).

In humans, papillary tumors grow over time into the eponymous shapes of long, finger-like protrusions with a fibrovascular core (Fig. 1J and Fig. S1C and D). In mice, the tumors do not form the elongated shapes seen in humans (Fig. 1K). One factor that potentially contributes to their absence in mice are the spatial restrictions in the much smaller mouse bladder. In the human bladder, the tumors have much more space to grow into the bladder lumen, and even a non-invasive pTa tumor, such as displayed in Fig. 1J, can be of the size, or even bigger than an entire mouse bladder. Nevertheless, in a biopsy from a noninvasive pTa tumor we found structures in the BM that have a striking similarity in size and shape to those seen in mice. (Fig. 1L and M). Therefore, we reason that the structures described above represent the early onset of papillary BC.

Thickness of the urothelium increases twofold in BBN-treated mice BC starts from a thickening of the urothelium in the form of hyperplasia or dysplasia [5]. Accordingly, we sought to quantify the thickness of the urothelium in the bladders. To this end, we implemented an image analysis pipeline which allowed us to quantify and visualize the thickness of the bladder urothelium in mice (Fig. 2A) as detailed in the Methods section.

**Figure 2:**
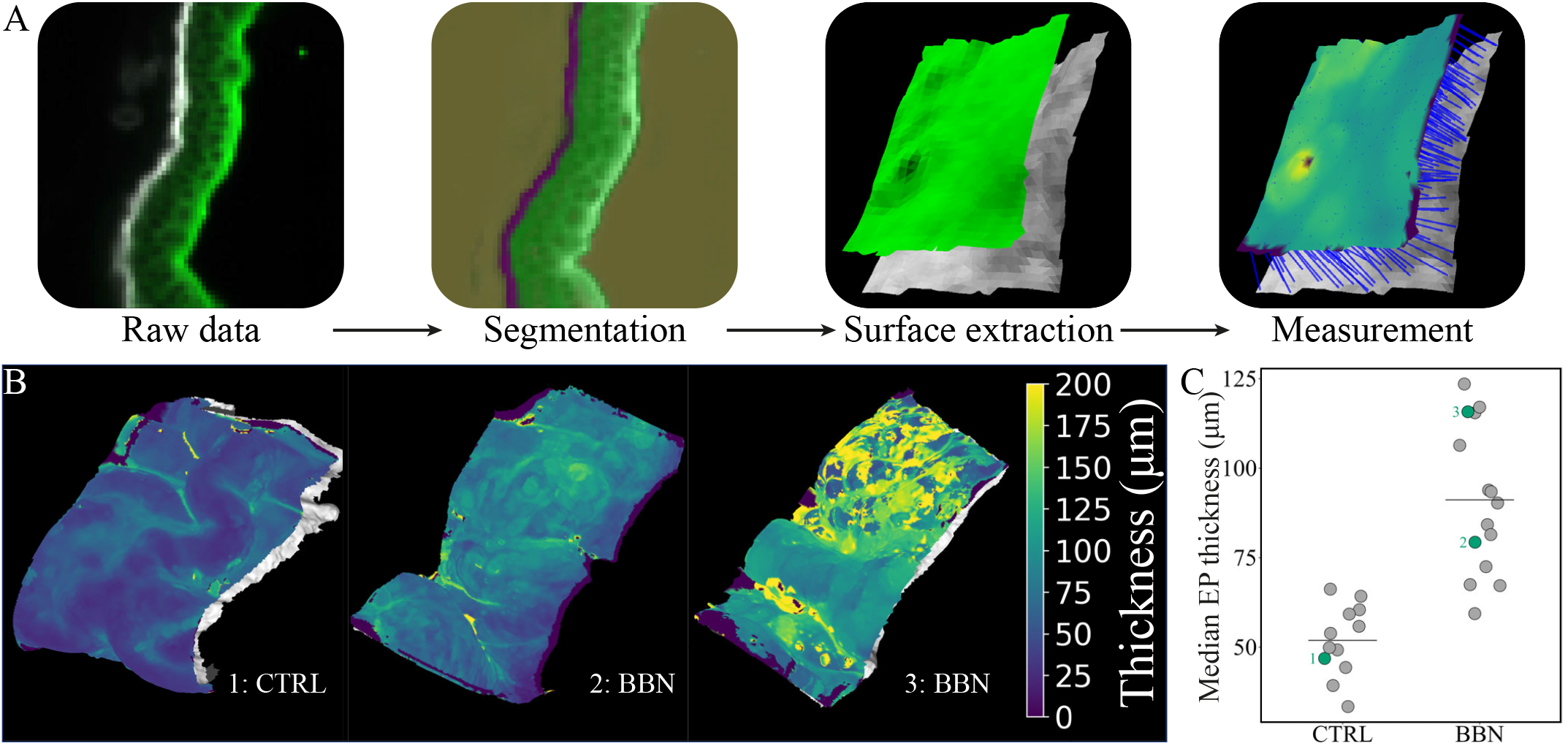
Thickness quantification of the urothelium. (A) Schematic representation of quantification pipeline used for thickness quantification, (B) Comparison of the urothelial thickness in control cohort mice and BBN treated mice 4 weeks post BBN, (C) Quantification of the thickness increase shows an almost two-fold increase in the thickness and a higher variance in the tissues from the BBN cohort. The green highlighted points correspond to the examples in (B).

Examples of the thickness measurement visualization are displayed in Fig. 2B. The left image is from a mouse bladder biopsy of a control mouse with a relatively homogenous thickness and a folding pattern exemplary for an empty bladder. The middle and right images show various degrees of thickening of the urothelium in mice 4 weeks after BBN treatment. The middle example shows a thickening of the urothelium in an area where the BM does not display extensive buckling (Fig. S2), while the right image is an example for the thickening of the urothelium in an area where clearly visible BM buckling is present (Fig. S2). In agreement with previous reports, we observe a substantial, almost twofold increase in the mean urothelial thickness of BBN-treated mice (91.2*±* 20.6 μm, SD) compared to controls (51.9*±* 10.0 μm, SD). Moreover, we noticed a higher degree of urothelial thickness variation within a single bladder, as well as between bladders in BBN-treated compared to control mice (Fig. 2C, S2).

### Mechanical simulations recapitulate different basement membrane morphologies

Thin elastic sheets and composites are known to exhibit a wealth of mechanical deformation modes under differential growth conditions or compression [39–45]. At large strains, secondary buckling modes can give rise to localized folding patterns [46, 47] reminiscent of papillae. To explore whether the formation of the different BC morphologies could be of mechanical origin, we employed computational continuum modelling using the finite element method (FEM), as detailed in the methods section. We simulated the mechanics of planar 3D sections of the bladder mucosa, 600 ×600 μm in size, consisting of three tissue layers: a 40 μm thick EP, a 1 μm thick BM and a 100 μm thick LP (Fig. 3A). The layer thicknesses vary between species; the human urothelium is about 45–110 μm thick [48]. The thickness of 40 μm for the EP layer was chosen as we start from a flat conformation in our simulations, corresponding to the EP’s slightly stretched state in a non-empty mouse bladder. By expanding the urothelium and the BM by up to 50% in both horizontal directions but restricting lateral outgrowth at the edges, we simulated local tissue overgrowth confined by the adjacent, healthy tissue. Assuming incompressible, isotropic linear elasticity in all three layers for simplicity, the Young’s moduli EEP, EBM, ELP describe the stiffness of each layer up to a plastic yield point (εy, σy), beyond which we included structural rearrangements on the cellular or subcellular scale into our model by making the EP and LP deform plastically (Fig. 3B, Methods).

**Figure 3:**
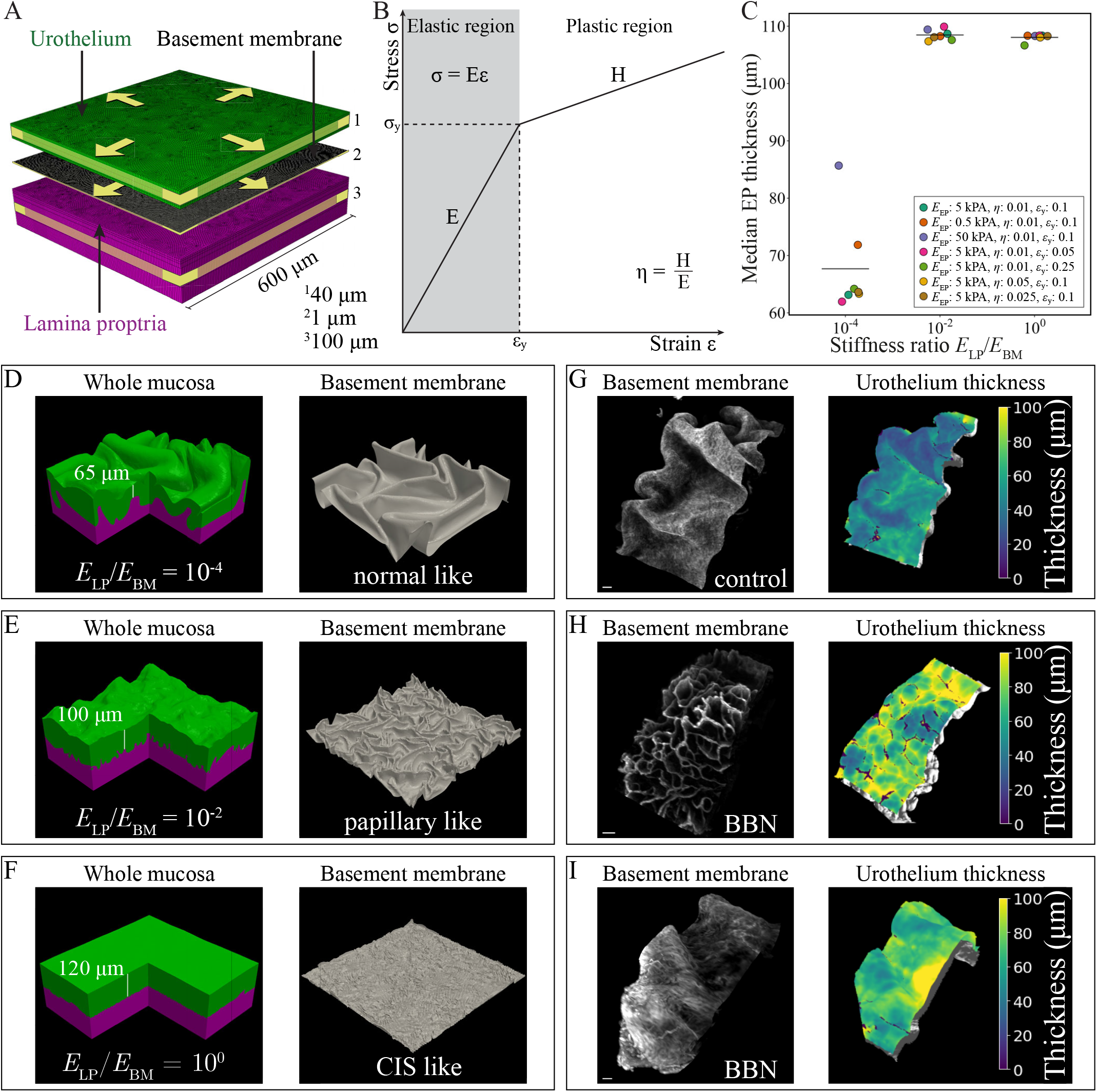
Different deformation modes in the BM depend on the stiffness of the LP and BM. (A) Schematic of the mechanical model setup, with volumetric expansion (yellow arrows) and boundary confinement (yellow bands), (B) Employed linear elasto-plastic constitutive stress-strain relationship, (C) Numerically observed thickening of the EP as a function of the stiffness ratio of the LP and BM, detailed parameter description can be found in Tab. S1, (D–F) Simulated mucosa morphologies at 50% expansion of the EP and BM in both in-plane directions, at indicated stiffness ratios, and Corresponding BM shapes, (G–I) Lightsheet microscopy images of the mouse bladder BM 4 weeks post BBN, showing normal smooth folding (G) and aberrant microscopic structures (H and I), and corresponding EP thickness profiles, scale bar: 100 μm.

The mechanical patterning is governed by relative stiffness ratios and layer thicknesses (see Supplementary Information). In numerical simulations at different plastic hardening moduli and yield points, we found that a decreasing stiffness ratio ELP/EBM affects the extent of thickening of the urothelial layer (Fig. 3C), with a mean thickness of 67.7 ± 8.6 μm (SD) for *E*_LP_/*E*_BM_ = 10^−4^, 108.5± 0.9 μm (SD) for E_LP_/*E*_BM_ = 10^−2^ and 108.0± 0.6 μm (SD) for *E*_LP_/*E*_BM_ = 1, and leads to the emergence of distinct folding patterns in the BM (Fig. 3D, E and F). Increasing or decreasing the stiffness of the urothelium *E*_EP_ results in thickening of this layer at *E*_LP_/*E*_BM_ ≈10^*−*4^, while only marginally affecting the folding patterns (Fig. S3). If *E*_LP_/*E*_BM_ decreases (because the BM becomes stiffer or the LP softer), the BM folding pattern coarsens. At *E*_LP_/*E*_BM_ ≈10^−4^, the morphology qualitatively resembles the normal smooth folding of the BM in an empty, healthy bladder (Fig. 3G). The undulations and folds transcend the BM and affect the entire urothelium, which thickens only little, similar to a normal urothelium in mice (Fig. 3G). At larger values of *E*_LP_/*E*_BM_, the folding occurs over shorter length scales, closer to the size of individual cells. At *E*_LP_/*E*_BM_ ≈10^−2^, sharp folds with small amplitude dominate the pattern, but translate only into mesoscopic bumps on the apical surface of the urothelium (Fig. 3F). The BM morphology qualitatively resembles that of BBN-treated mice, which can show networks of mesoscopic, papillary-like creases, and whose EP exhibits elevated thickness (Fig. 3H). At LP and BM stiffness parity, undulations on the apical surface of the EP disappear, and the majority of its volumetric expansion is translated into thickening. The BM exhibits microscopic folding and crumpling (Fig. 3F) akin to the fine-grained, non-uniform (CIS-like) structure observed atop the macroscopic folds in BBN-treated mice (Fig. 3I). The EP thickness profile in simulations agrees with the observed, elevated but more uniform one observed in imaging (Fig. 3I).

### Stiffness ratios govern mucosa folding modes

To obtain a more comprehensive, quantitative insight into the mucosa morphologies, and to further examine the potential of tissue mechanics to disrupt its structural integrity at the onset of BC formation, we sought a geometrical quantity that characterizes the emergent BM structure. We extracted the simulated BM height maps (Fig. 4A) and Fourier-transformed them (Fig. 4B) to determine the geometrical spectrum of wrinkles and folds (Fig. 4C, see methods for details). The dominant wave-length *λ*_0_ encodes the primary spatial separation between creases and furrows in the BM. Fixing all other model parameters, we computationally screened the space of mucosa deformations spanned by the stiffness ratios between the three layers, considering a substantially stiffer BM than the EP and LP [28]. This analysis revealed a rich morphological phase space with two distinct regimes (Fig. 4D). At large values of E_EP_/E_BM_ but small *E*_LP_/*E*_BM_, the comparably stiff expanding urothelium buckles and folds along with the BM, a regime we term “plate-like” EP deformation, characterized by undulations with a large wavelength in the order of ten cell diameters or more. Conversely, at comparably stiffer LP than EP (large values of *E*_LP_/*E*_BM_ but small *E*_EP_/*E*_BM_), it is the BM alone that buckles and folds, with the EP and LP effectively acting as elastic environment. In this “medium-like” EP deformation regime, disordered folds are observed in the BM with short dominant wavelengths in the order of only a few cell diameters or even below a single cell diameter. An energy balance based on linear elasticity theory [49, 50] yields a morphological phase boundary between the two regimes (Fig. 4D, black dashed line; see Supplementary Information for details), roughly following the *λ*_0_ ≈ 100 μm isoline. Traversing this boundary involves a steep change in the folding wavelength (Fig. 4D, colored contours), as it is proportional to h_EP_ in plate-like buckling, and proportional to *h*_BM_ in medium-like buckling (Supplementary Information). At fixed epithelial stiffness, the shape transition occurs along the *E*_LP_/*E*_BM_ axis (Fig. 4E) with tighter folding toward larger values of *E*_LP_/*E*_BM_. An overview of all the different patterns observed in the simulations for Fig. 4D and E can be found in Fig. S4A and S5; the median thickness of a subset of simulations from Fig. 4D in Fig. S4B.

**Figure 4:**
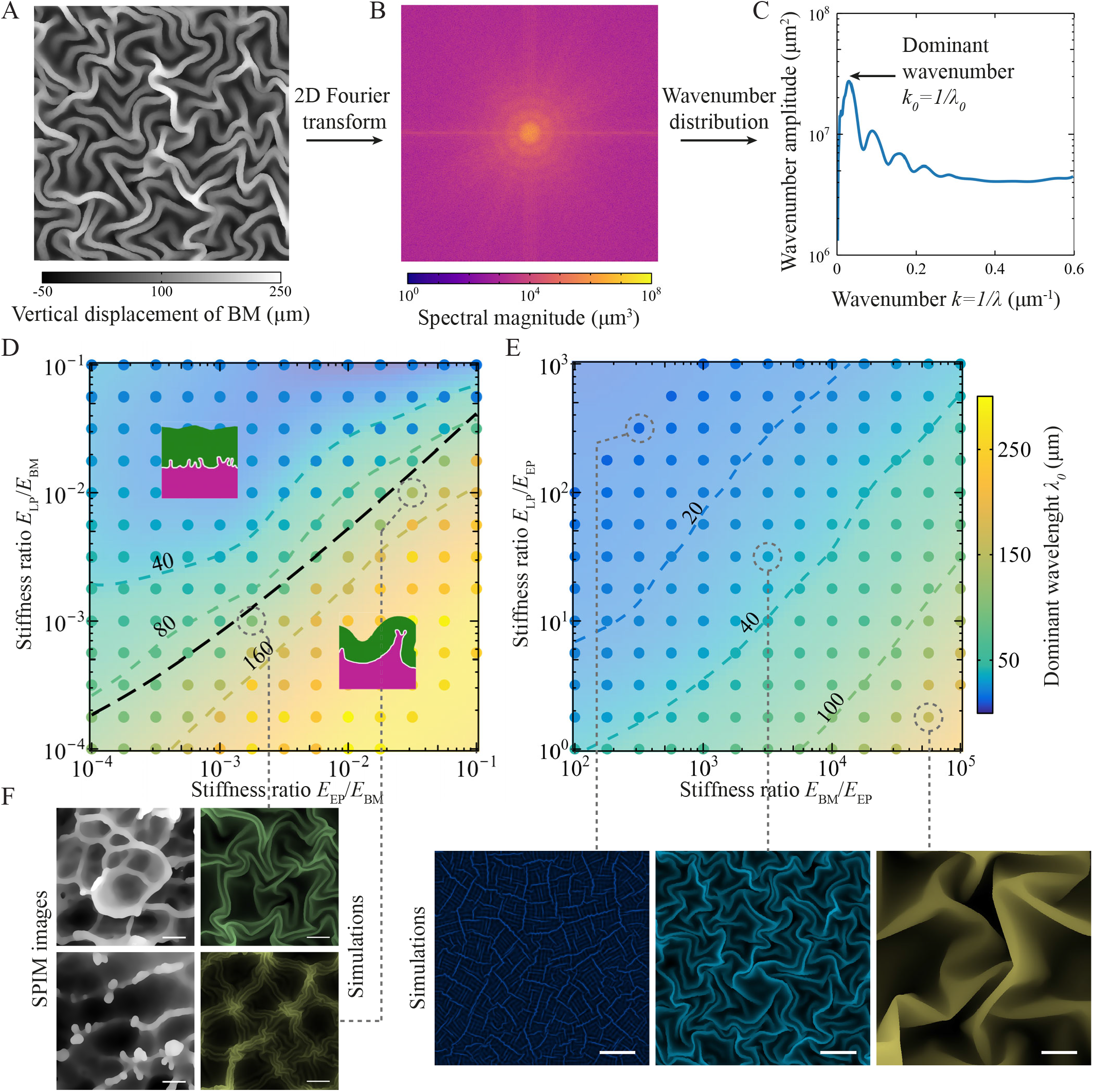
Morphogenetic analysis of simulated BM buckling patterns. (A) Exemplary height map of the BM showing overhanging folds. (B) Spatial Fourier transform of (A). (C) Wavenumber distribution calculated from a phase average of the spectral magnitude. (D–E) Numerically obtained morphological phase diagram as a function of tissue stiffness ratios. Dots represent dominant wavelengths of individual simulations. Inset images show exemplary vertical slices of two deformation regimes, separated by a theoretical boundary (black dashed line). See Supplementary Information for details. Simulation parameters: *h*_EP_ = 40 μm, *h*_BM_ = 1 μm, *h*_LP_ = 100 μm, *η*_EP_ = 0.01, *η*_LP_ = 0.01, *ϵ*_y,EP_ = 0.1, *ϵ*_y,LP_ = 0.1. (F) Examples of vertical BM displacement from SPIM images juxtaposed to simulations at marked stiffness ratios (dashed gray lines). Simulation images are colored according to their dominant wavelength, scale bar: 100 μm.

Plausible locations of the mouse and human bladder mucosa in the stiffness plane range from a vicinity of the morphological boundary toward greater E_LP_/E_EP_ ratios. With tangent moduli of the urothelium in the order of 2–6 kPa [51] and about two to three orders of magnitude larger in the BM [52], the physiological region lies in the middle to upper part of Fig. 4D, where the folding pattern is sensitive to small stiffness changes.

Comparing numerical simulations with SPIM images of the mouse BM, we found similar geometrical features, although the numerical model rests on simplifying assumptions and does not reflect the biological inhomogeneity present in real tissues. Simulations close to the morphological boundary yield troughs surrounded by ridges in the BM (Fig. 4F, top row) at *E*_EP_/*E*_BM_ ≈*E*_LP_/*E*_BM_≈ 10^−3^. For stiffer EP and LP (or softer BM), spot-like stress condensates are observed, with ridges connecting them at varying levels of BM elevation (Fig. 4F, bottom row)—a feature reminiscent of the onset of the finger-like protrusions in papillary tumor formation. Strikingly, these protrusions emerge facing the bladder lumen in the simulations like they do in papillary tumors.

### Basement membrane softens locally in BBN treated mice

Our numerical simulations demonstrate that the buckling pattern can be explained by stiffness changes in the different layers of the mucosa. Softening of urothelial cells [53–55] and the upregulation of different ECM-related genes, linked to stiffening of the LP [37, 56, 57], have been investigated in BC and recognised as a risk factor for progression and invasion in BC. We were interested in investigating whether BC would also cause a change in stiffness in the BM. We used AFM-based indentation measurement on decellularised mucosas from BBN-treated mice 4 weeks after treatment and corresponding mucosas from the control cohort (Fig. 5A). The mean overall stiffness of the BM was 120 kPa in the control cohort and 97 kPa in the BBN cohort (Fig. 5B, Tab. S2), indicating that the BM is on average softer in the BBN cohort. Furthermore, if the different measurement positions are examined separately, some measured spots in the BBN cohort display a similar BM stiffness as the control cohort, while some areas show an almost 6-fold softening of the BM compared to the overall stiffness in the control group (Fig. 5B, Tab. S2). This confirms that BM softening does indeed occur early during BBN treatment in mice.

**Figure 5:**
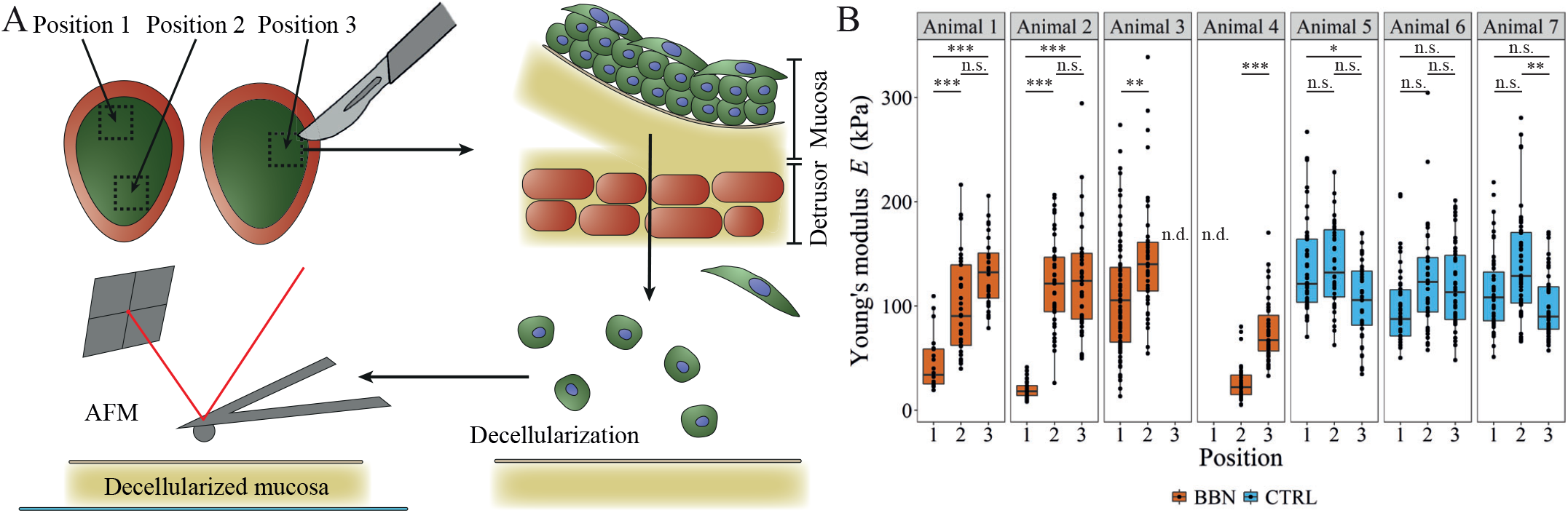
BM shows localised softening in AFM. (A) Visualization of the sample processing for AFM. (B) BM stiffness at 2-3 positions in bladders from the BBN treated cohort 4 weeks post BBN and corresponding control cohort mice, with adjusted P values. Significance levels: *P* ≤ 0.001, ***; *P* ≤ 0.01, **; *P* ≤ 0.05; *P* > 0.05, n.s.

## Discussion

Based on our morphometric imaging analysis, we propose that a mechanical buckling instability underlies the distinction between different bladder carcinoma subtypes. Our mechanical computer simulations revealed a morphological transition in the mucosa depending on the relative stiffnesses of its constituent layers, leading to distinct buckling patterns upon overgrowth of the urothelial layer. Intriguingly, the physiological regime lies in the morphological transition zone where structural changes require only small stiffness changes, hinting at a delicate constitutive balance between the EP, BM and LP. Perturbations of this balance accompanied by differential growth can lead to epithelial thickening and BM folding with short wavelengths. Our simulations and theory show that these morphological changes are strongest approximately along the *E*_LP_/*E*_EP_ axis. This suggests two mechanical mechanisms that may disrupt the structural integrity of the bladder mucosa at the onset of BC formation: Alterations in the urothelial-to-membrane stiffness ratio *E*_EP_/*E*_BM_ or in the laminal-to-membrane stiffness ratio *E*_LP_/*E*_BM_ can both give rise to a shortening in the BM folding lengthscale, potentially down to the (sub)cellular level, where mechanical tissue damage may ensue. We found striking morphological agreement between the BM structure observed in simulations and in mice with developing BC, induced by BBN, providing a mechanical candidate explanation for the emergence of papillary tumors and CIS.

Our morphometric imaging analysis revealed that almost identical structures of the BM as found in the mouse, can also be observed in humans. This suggests that papillary tumors in humans and mice develop through similar mechanical means during early tumor growth. Such mechanical instabilities have previously been explored only in the context of tumor spheroids [58, 59]. Generally, solid mechanics models (for instance, coupling ECM mechanics to cellular behavior) remain underrepresented in cancer research [60, 61].

Our results echo with recent multilayered 2D vertex model simulations that found that stiffening of the BM promotes folding in skin cancer [7]. Our findings embed this result in a greater perspective of a mechanical patterning mechanism that explains not just single folds, but entire lesions. They also align with studies that have demonstrated that BC cells from more aggressive origin are softer than cells from less aggressive BC origin [53–55]. The upregulation of type I and III collagen genes, predominantly found in the LP [62], is considered a marker for increased ECM stiffness and linked to BC invasion [37]. On the other hand, overexpression of the collagen IV proteases MMP2 and MMP9 have been recognized as a marker for BC progression [63]. Collagen IV is, besides Laminin, one of the main ECM proteins of the BM [62]. The degradation of collagen IV could explain why we observe localized softening of the BM in BBN mice. Both the stiffening of the LP and the softening of the urothelium and BM are changes that our model predicts to support lesion formation.

Our computational model is necessarily a simplification of the real mucosa and has limitations. For example, we do not take the vascularisation of the LP into account, despite the known role of angiogenetic processes in many cancers, including BC [64]. In our SPIM images, we can observe that the vasculature always follows the buckling pattern and undulations of the BM. We hypothesise that for the formation of elongated and branched structures, as seen in human papillary tumors, vascularisation also acts as an additional driver or structural scaffold. Moreover, we assumed isotropic linear elasticity in all mucosa layers (with plastic strain hardening in the urothelium and LP to incorporate stress and strain localisation due to the tissue’s ability to relax stress with cellular rearrangements). While this is likely reasonably accurate for the near-threshold buckling behavior at the lesion onset, more elaborate constitutive behavior may be appropriate for large deformations of the bladder [65]. Experimental measurements of these are challenging, but would be a valuable addition to the scientific pool of data, to be acquired in future research.

A recent study in rats found that BBN treatment leads to a softening of the LP in the BBN treatment cohort. However, this does not contradict our finding since, overall, the LP stiffens over the time course of the treatment due to the ageing of the rats [51]. In BC, the patient’s age is one of the risk factors for cancer development and progression [66], which could be reflected by the stiffening of the ECM due to ageing. Furthermore, urinary schistosomiasis [67] and previous radiotherapy of prostate cancer are known risk factors for developing BC [68, 69], both of which induce fibrosis in the bladder [67, 70], and recently, tissue stiffening due to transurethral resection (TURB) was proposed as a risk factor for BC progression [71]. This suggests that the tumor-independent processes can also act as drivers of the stiffening of the LP in BC. While the stiffness changes in the different mucosa layers were previously mostly investigated in the context of cancer progression, we demonstrate here that these changes could play an important role already at the onset. Further research is required to test this prediction experimentally, and to get a clearer picture of the magnitude, direction, timing, and causation of changes in the mechanical properties of the mucosa layers, and to what degree these changes are induced by the cancer itself or by cancer-independent processes.

For low-risk non-muscle-invasive BC, treatment usually consists of TURB in combination with intravesical chemotherapy [17]and has a good 5-year survival prognosis [72, 73]. However, BC often reoccurs and has a probability of 0.8%–45% to progress to muscle-invasive BC [15]. Especially CIS have a high risk for progression compared to papillary tumors [16, 17]. Although treatment options for muscle-invasive BC and metastatic disease have been expanded to immunotherapy and targeted therapies in recent years [74], the prognosis for survival remains substantially less favourable [73].

Incorporating stiffness measurements into the diagnostic process, e.g., through shear wave elasticity examinations [57] or AFM on biopsies [75], could help to improve patient risk stratification and inform personalized treatment and monitoring regimes. Additionally, treatment advancements that prevent excessive scar formation may not only help to maintain normal bladder function and enhance the quality of life for patients [76] but might also lower the risk of BC recurrence and progression. In light of our study, the structural integrity and mechanical stability of the mucosa layers offers a promising line of attack for medical treatment and prophylaxis of BC progression.

## Methods

### Ethical Statement

Human BC biopsies were provided by the University Hospital of Basel (USB), Switzerland, under approval by the Ethical Committee of Northwestern and Central Switzerland (EKBB 37/13).

All experiments involving animals were performed in accordance with the Swiss animal welfare legislation and approved by the veterinary office of the Canton Basel-Stadt, Switzerland (approval number 2957/29841). All animals were housed at the D-BSSE/University Basel facility under standard water, chow, enrichment, and 12-h light/dark cycles. An overview of all animals used in this study can be found in Tab. S3.

### Mouse strain and bladder cancer induction

To distinguish the urothelium from the other tissue layers in the bladder we used the Shh^*Cre/+*^;Rosa26^*mT/mG*^x RjOrl:SWISS crosses (Shh^*Cre/+*^;Rosa26^*mT/mG*^) previously described by Conrad et al. [38]. In these mice, the fluorescent marker m-tdTomato is expressed through the bladder tissue except in the urothelial layer, which expressed mEGFP.

BC can be induced in mice by spiking the drinking water with N-Butyl-N-(4-hydroxybutyl) nitrosamine (BBN) [18]. BBN is known to induce an inflammatory response already in the first weeks of treatment and BC few weeks after BBN stop [77]. The ideal sampling time point for early stages of BC was found to be 4 weeks post BBN. 11 weeks post BBN was defined as the latest sampling point to avoid suffering due to excessive tumor growth. In our experimental setup, 10-week-old male mice were provided with 0.05% BBN (Sigma-Aldrich) in the drinking water for 12 weeks followed by 4–11 weeks normal drinking water. For the control condition, we used littermates maintained under identical housing conditions. To avoid suffering due to unwanted, excessive tumor growth mice were frequently monitored for signs of distress and body weight was controlled weekly.

Mice were sacrificed at desired time point post BBN, and the bladders immediately harvested and washed in cold DPBS (Gibco). Excess fatty tissue was removed with surgical scissors and forceps before cutting the bladders sagittally into halves. For imaging, the bladder halves were fixed in 4% paraformaldehyde (PFA) (Thermo Fisher) for 3–4 h at 4°C and subsequently processed as described below. Bladder halves used for AFM were submerged in Betadine solution (povidone-iodine 11 mg/mL, Mundi Pharma) for 1 min and rinsed with DPBS. Further processing is described under the subsection AFM.

### Optical clearing and immune fluorescence staining of fixed bladder tissue

Whole-mount tissue clearing of human and mouse biopsies was performed using the Clear Unobstructed Brain/Body Imaging Cocktails and Computational Analysis (CUBIC) protocol [78]. For better clearing and imaging, human and mouse biopsies were further cut into smaller sections if needed. Clearing times in reagents for decoloring, delipidation, permeation (CUBIC-1), and refractive index (RI) matching (CUBIC-2) were adjusted to maximize clearing efficiency and minimize quenching. Biopsies were incubated overnight in 50% CUBIC-1 (CUBIC-1:H2O, v/v) at 37°C, followed by incubation in 100% CUBIC-1 for 6-10 days at 37°C on a nutating shaker. After optical clearing, the biopsies were washed three times in PBS on a rotating mixer for > 1 h at room temperature (RT). All reagents used for preparation of CUBIC-1 and CUBIC-2 were obtained from Sigma-Aldrich, with the exception of N,N,N’,N’-Tetrakis(2-hydroxypropyl)ethylenediamine (Tetrakis) obtained from TCI and Polyethylene glycol (PEG) mono-p-isooctylphenyl ether (Triton X-100) obtained from Nacalai Tesque (discontinued).

For immunofluorescent labelling, cleared biopsies were blocked for 3–4 h at room temperature (RT) or overnight at 4°C in blocking buffer (PBS (Sigma-Aldrich), 10% Fetal Bovine Serum (FBS) (Sigma-Aldrich), 1% Bovine Serum Albumin (BSA) (Sigma-Aldrich), 0.2% Triton X (Sigma-Aldrich) and 0.02% sodium azide (Sigma-Aldrich)). Blocked samples were then incubated with primary antibodies in blocking buffer for two nights at 4°C. After incubation with primary antibodies, samples were washed again with PBS three times for > 1 h at RT on a rotating mixer and subsequently incubated again for two nights at 4°C with secondary antibodies in blocking buffer. For mouse tissues chicken anti-GFP (Aves Labs; GFP-1020; 1:500) and rabbit anti-laminin (Abcam; ab11575; 1:500) primary antibodies, and for human biopsies (see Fig. S1B and C) goat anti-ZO-1 (Thermo Fisher; PA5-19090; 1:200) and anti-laminin primary antibodies were used. As secondary antibodies the following fluorescently labelled antibodies were used: Alexa Fluor 488 goat anti-chicken IgG (Invitrogen; A11039; 1:500) and Alexa Fluor 647 donkey anti-rabbit IgG (Invitrogen; 32795; 1:500) for mice, and Alexa Fluor 488 donkey anti-goat IgG (Invitrogen; 11055; 1:500) and Alexa Fluor 647 donkey anti-rabbit IgG for human biopsies.

At the end of the incubation, the samples were briefly washed in PBS before post-fixation for 15min in 4% PFA at RT. To wash off the remaining PFA, the samples were washed three times for > 15 min at RT in PBS before being placed in 50% CUBIC-2 (CUBIC-2:H2O, v/v) overnight at RT, followed by 6–10 days in 100% CUBIC-2.

### Embedding and imaging cleared samples

Cleared samples were embedded in 2% low-melting-point solid agarose cylinders and first immersed in 50% CUBIC-2 for one night and then in 100% CUBIC-2 for up to two days. 3D image stacks were acquired using a Zeiss Lightsheet Z.1 SPIM using a 5x/1.0 clearing objective. For large samples, tiled acquisition was used to image the whole sample. For each sample, laser powers and exposure times were adapted to the signal intensity to ensure optimal image quality for all acquisitions.

### Image Processing and Analysis

Tiled images were reconstructed using the Fiji [79] plugin BigStitcher [80]. For 3D visualization we used the software Imaris (Oxford Instruments).

For thickness quantification, single tiles were selected and scaled in Fiji, such that the *xy* pixel size matches the pixel size in *z* direction. To extract surface boundaries, we first performed pixel classification using the machine learning tool ilastik [81] to create tissue predictions for the urothelium and the BM. These predictions were put into a custom Python script to generate triangulated meshes. The meshed were subsequently post-processed and cleaned using MeshLab [82] and the integrated Screened Poisson Surface Reconstruction algorithm [83].

To approximate the thickness of the urothelium, we used a normal ray-based approach by projecting normal rays from the urothelium-lumen interface towards the urothelium-BM interface. The length of the rays to the point where they intersect with the urothelium-BM interface was then used to approximate the tissue-thickness of the given position (Fig. 2A). The surface meshes of the urothelium, with a color map representing the local thickness, and the BM were visualized using Napari [84]. The Python scripts are available as Jupyter notebooks (see Code Availability).

### Numerical Simulations

We simulated the nonlinear mechanical deformation of the mucosa in response to differential volumetric growth using the finite element method in Abaqus FEA 2021 (Dassault Systèmes). Model files are provided (see Code Availability). The mucosa was represented as a three-layered continuum consisting of the EP, the BM and the LP, from top to bottom. We simulated square tissue sections with an edge length of 600 μm, and initial layer thicknesses *h*_EP_ = 40 μm, *h*_BM_ = 1 μm and *h*_LP_ = 100 μm (Fig. 3A). Boundary conditions were set to constrain the vertical displacement of the bottom surface of the LP and the horizontal normal displacement on all four lateral sides of all three layers. To mimic epithelial proliferation, we expanded the volumes of the EP and BM layers orthotropically by 50% in each of their planar directions, but not vertically, resulting in a total volumetric fold-change of 2.25 in these two layers. In response to the stresses caused by this differential growth and the confined tissue boundaries, the mucosa buckled out of plane.

In absence of detailed knowledge on the exact constitutive relationships in healthy and cancerous mucosa, we assumed isotropic, homogeneous, linearly elasto-plastic behavior in all three layers. For a more detailed discussion of constitutive bladder models, see [65]. The stiffness in the elastic deformation of each layer was set by their Young’s moduli *E*_EP_, *E*_BM_ and *E*_LP_. To make the mucosa nearly incompressible, we fixed all Poisson ratios at *v* = 0.48. Local stress relaxation at the cellular or sub-cellular level was incorporated in our model by making the EP and LP plastic upon reaching a von Mises stress yield stress *σ*_y_ = *Eϵ*_y_, where *ϵ*_y_ is an effective yield strain (Fig. 3B). We fixed *ϵ*_y_ = 0.1 in both the EP and LP, a value that can be quickly reached as the EP and BM expand by 50% in both of their planar directions. In the plastic regime, linear strain hardening was assumed for simplicity, with isotropic hardening moduli *H*_EP_ and *H*_LP_. We kept the ratios *H*_EP_/*E*_EP_ = *H*_LP_/*E*_LP_ = 0.01 fixed in all simulations, except where specified otherwise.

The EP and LP were modeled as solids, discretized into 125,111 and 142,984 hexahedra (element type C3D8R), respectively. The BM was modeled as a thin shell consisting of 78,239 triangles (element type S3R). We explicitly solved Newton’s dampened second law, making sure that inertial and viscous forces remained small throughout.

### Analysis of the wrinkling pattern

Upon complete volumetric expansion, we extracted vertical displacement maps *w*(*x, y*) of the BM in parallel projections as seen from the top (Fig. 4A), at a resolution of 1.2 pixel/μm. As our simulations were geometrically fully nonlinear, the simulated BMs can overhang (Fig. 4D, insets), just like in the lightsheet images. To quantify the tissue morphologies, we performed a spectral analysis on the height maps using the 2D Fourier transform (Fig. 4B),

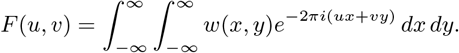

*F*(*u, v*) is generally complex, and the spectral magnitude

|*F*(*u, v*)| encodes periodic undulations in the BM with wave vector 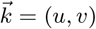. To extract the isotropic (radial) component in the wrinkling pattern, we averaged over all polar angles:

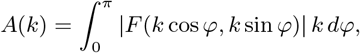

where 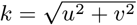 is the wavenumber. This amplitude can be interpreted as a measure of isotropic spatial autocorrelation in the vertical displacement field of the BM. It typically decays for increasing wavenumbers, exhibiting local maxima at the predominant wavelengths *λ*_*i*_ = 1/*k*_*i*_, *i* = 0, 1, … (Fig. 4C). The wavelength with largest amplitude, *λ*_0_, quantifies the primary buckling mode, and served as a scalar measure of the main average spatial separation between wrinkles or folds in the mucosa (Fig. 4D and E).

### AFM-based indentation measurement

Bladder halves were further cut into approximately 3 ×3 mm specimens. Using fine forceps, the mucosa was carefully peeled off the detrusor. From each of the three control bladders, we selected three mucosa specimens from different locations of the bladder. Since the mucosa seems to be more fragile in BBN-treated mice, we were only able to obtain 2 mucosa specimens from two of the four mice from the BBN-treatment cohort. From the remaining two bladders, three mucosa specimens were obtained. For decellularization, the mucosas were submerged in a sterile 0.1% SDS (Sigma-Aldrich) solution and incubated at RT for 72 h on a rotating mixer. The SDS solution was refreshed every 24 h. After that, the mucosas were washed in sterile water for 72 h at RT on a rotating mixer and the water was replaced every 24 h. The decellularized mucosas were placed on a 35 mm glass bottom petri dish (World Precission Instruments) with the BM facing upwards. To improve adhesion, excess water was left to evaporate 5 min, so that the mucosas stick to the petri dish, before the dish was filled with sterile water.

An AFM (NanoWizzardII, JPK, Germany) was mounted onto an inverted light microscope (Observer Z1, Zeiss, Germany). A tipless triangular cantilever (NPO-D, Bruker, US) with a 5 μm diameter bead attached to its apex was used for the measurements. The spring constant of the cantilever was determined by the thermal noise method. For indentation experiments, the beaded cantilever was positioned above an area of interest. A grid of 4× 4 positions within 100 ×100 μm was defined for indentation experiments. The beaded cantilever was approached to the sample at 5 μm/s until a force of 10 nN was recorded. Subsequently, the cantilever was retracted and moved to the next position in the grid. Three different regions per sample were tested if possible. Data analysis was performed with the inbuilt JPK Data Analysis software. The recorded force-distance curves of the approach were offset and tilt corrected and corrected for the cantilever deflection. Afterwards, the approach force-distance curve was fitted by the Hertz model with a Poisson ratio of *v* = 0.5. The contact point in FD curves was determined automatically.

## Code Availability

The source code is publicly released under the 3-clause BSD license as a git repository at https://git.bsse.ethz.ch/iber/Publications/2023_lampart_bladder_cancer.

## Acknowledgements

This work was funded by PHRT iDoc grant 306 and SNF Sinergia grant CRSII5_170930. The authors thank Radhe Kumar for her contributions as part of her student projects, the imaging facility of the D-BSSE for training and technical support and the members of the Iber group for valuable discussions.

## Author Contributions

D.I., F.L.L., R.V., C.L.M, L.B., C.A.R., H-H.S., and D.J.M. designed the research; F.L.L., M-D.H. and G.C. performed animal experiments; C.A.R. provided human tumor biopsies; L.B. performed tumor staging and grading on human and mouse tumors; F.L.L. generated all imaging data; F.L.L. and K.A.Y analyzed the imaging data; R.V., F.L.L., Y.W., F.M. and D.I. developed, simulated, and analysed the computational model; N.S. and F.L.L. did the AFM measurements; F.L.L., R.V., K.A.Y, N.S. and D.I wrote the manuscript; F.L.L., R.V. and Y.W. created the figures.

