## Supplementary_Data for "Morphometry and mechanical instability at the onset of epithelial bladder cancer"

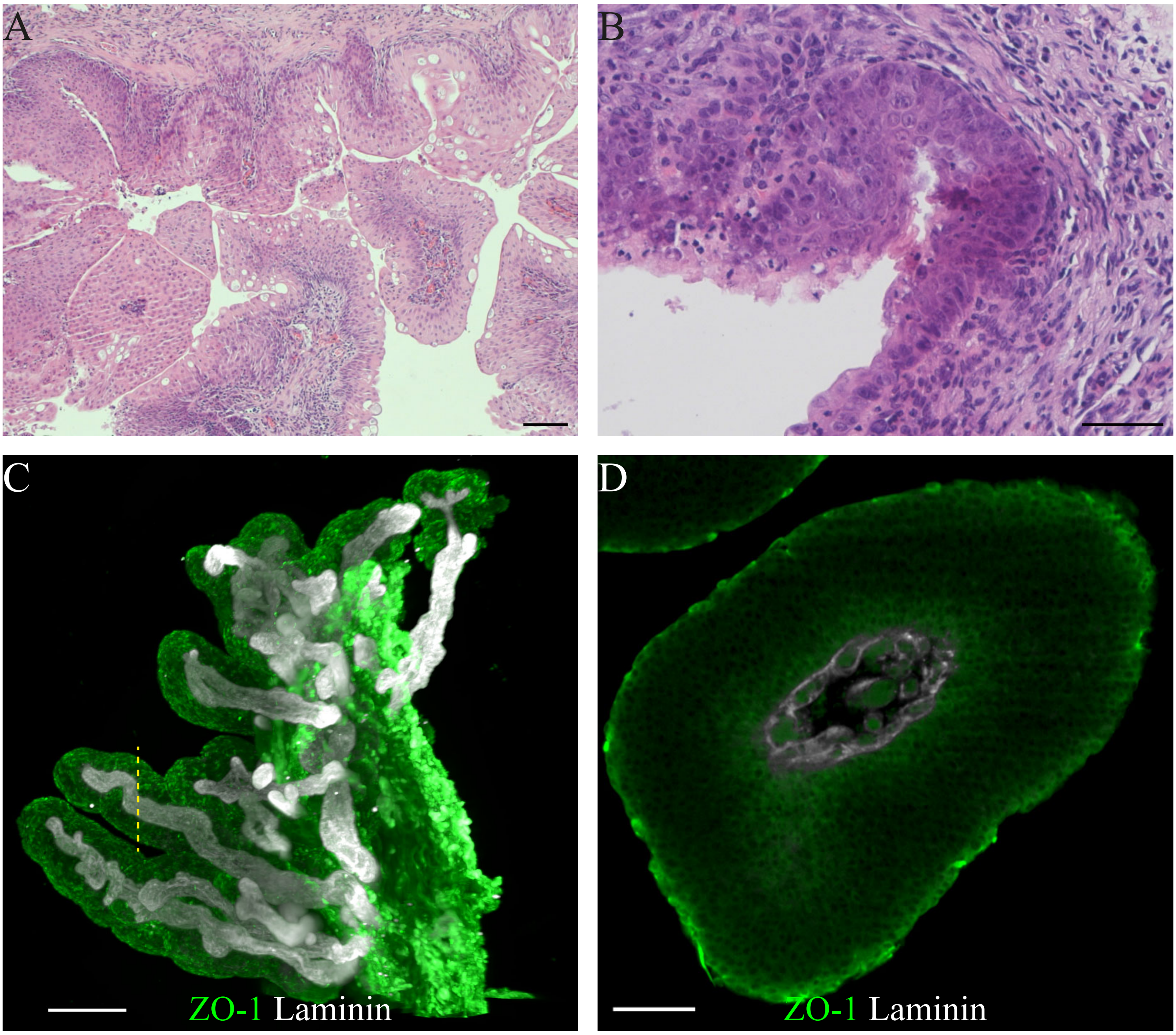


**Figure S1: Bladder tumors in mice and humans** (A) Low grade papillary tumor in the mouse bladder 11 weeks post BBN, scale bar: 100 µm, (B) CIS in the mouse bladder 8 weeks post BBN, scale bar: 50 µm, (C) Human papillary tumor, yellow dotted line: approximate position of (D), scale bar: 500 µm, (D) (digital) cut thru human papilla, scale bar: 100 µm.

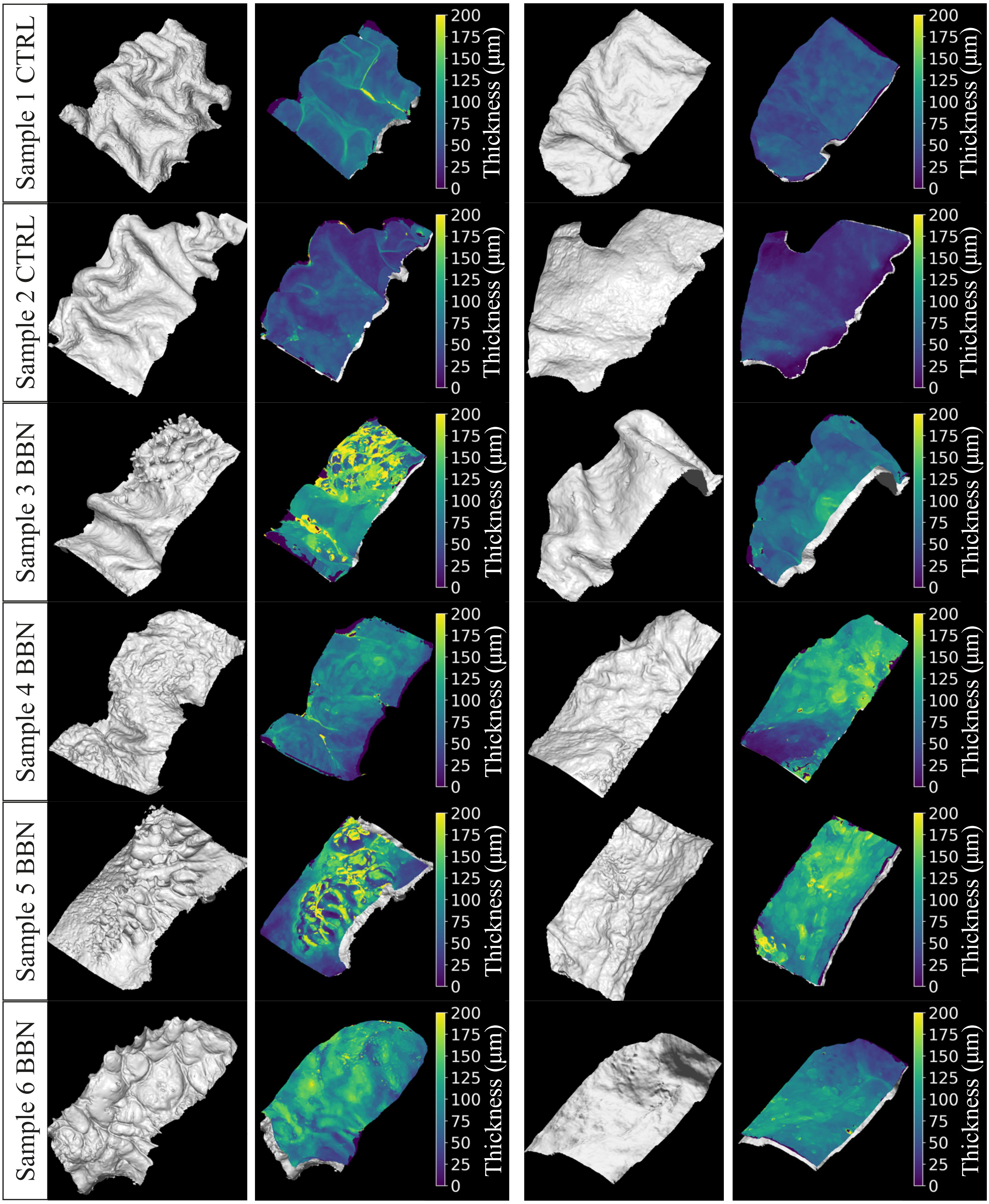


**Figure S2: Examples of different thickness measurements in mice** Comparison of different degrees of BM alterations and thickening in mouse biopsies from six different bladders, two tissue samples from each bladder.

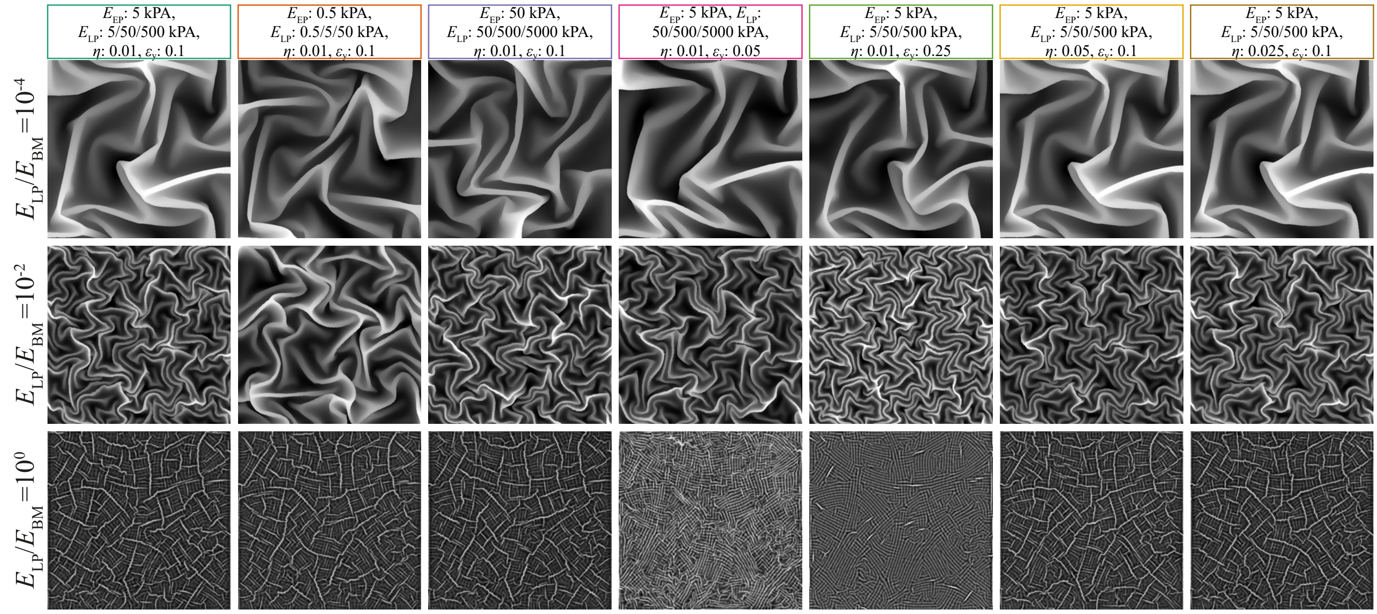


**Figure S3: BM shapes** Overview of all observed BM height profiles from the simulations in Fig. 3C with individually normalized contrast. The box colors correspond to the colors of the data points in the plot.

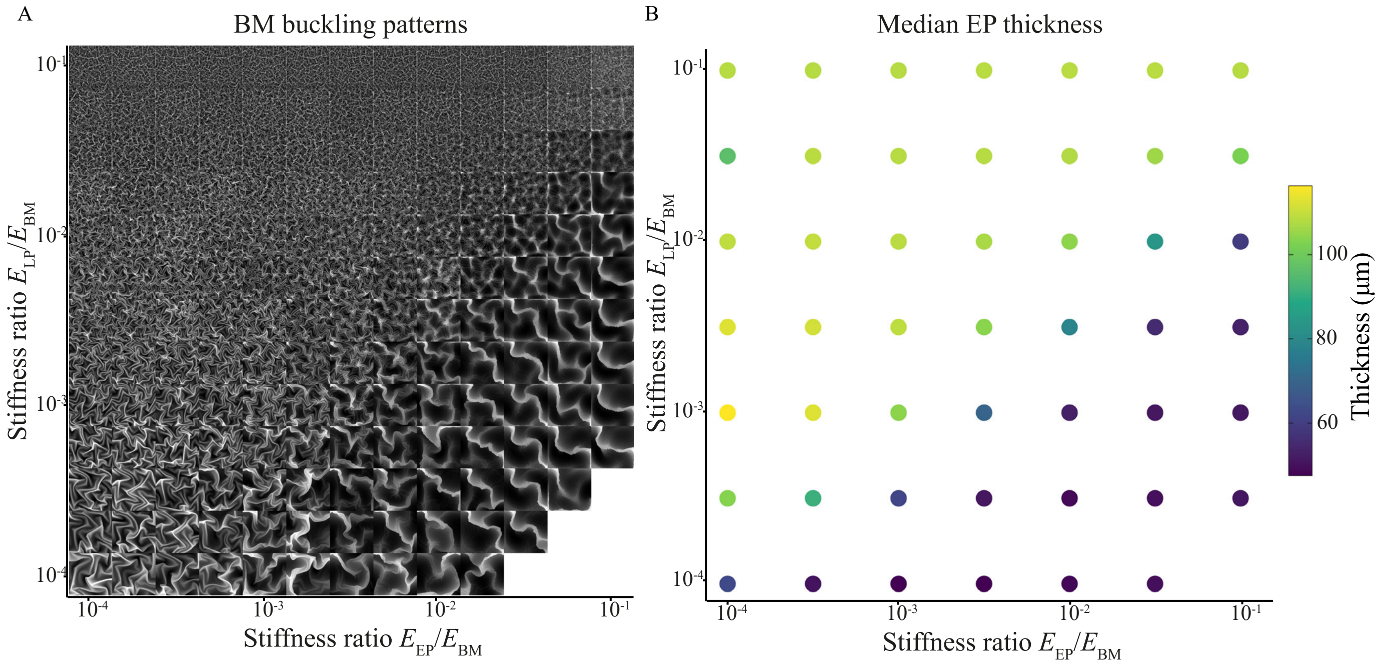


**Figure S4: BM shapes and EP thickness** (A) Overview of all observed BM height profiles from the simulations in Fig. 4D with individually normalized contrast, (B) Overview of a subset of thickness measurements from the simulations in Fig. 4D.

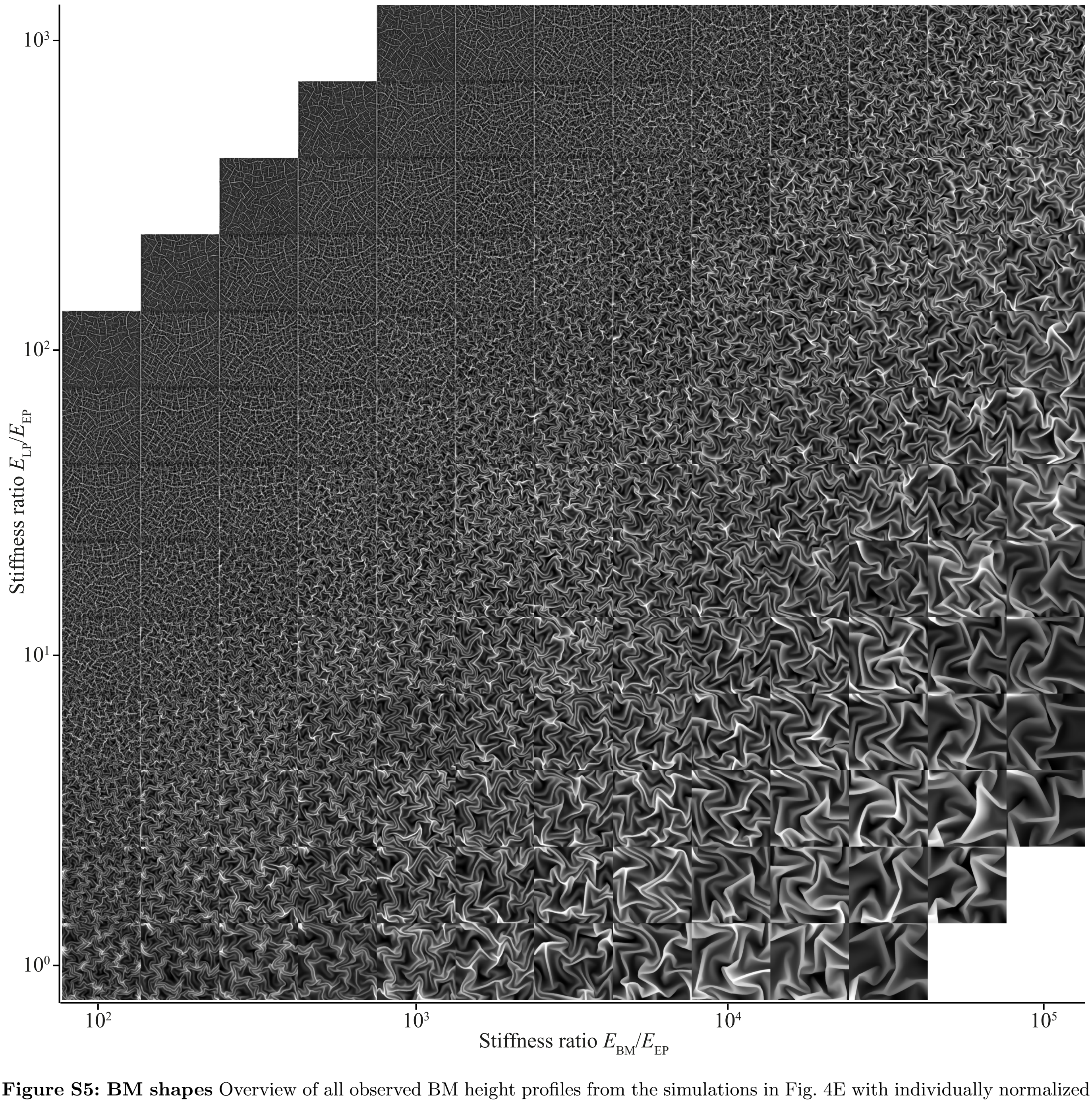


**Figure S5: BM shapes** Overview of all observed BM height profiles from the simulations in Fig. 4E with individually normalized contrast.

**Table S1: Simulation parameters** Overview of all parameters from the simulations displayed in Fig. 3C.

| | | Young's modulus $E$ (kPa) | | | Hardening $\eta$ | | Yield strain $\epsilon_y$ | |
|---|---|---|---|---|---|---|---|---|
| Simulation | $E_{LP}/E_{BM}$ | EP | LP | BM | EP | LP | EP | LP |
| 1 | $10^{-4}$ | 5 | 5 | 50000 | 0.01 | 0.01 | 0.1 | 0.1 |
| | $10^{-2}$ | 5 | 50 | 5000 | 0.01 | 0.01 | 0.1 | 0.1 |
| | $10^{0}$ | 5 | 500 | 500 | 0.01 | 0.01 | 0.1 | 0.1 |
| 2 | $10^{-4}$ | 0.5 | 0.5 | 5000 | 0.01 | 0.01 | 0.1 | 0.1 |
| | $10^{-2}$ | 0.5 | 5 | 500 | 0.01 | 0.01 | 0.1 | 0.1 |
| | $10^{0}$ | 0.5 | 50 | 50 | 0.01 | 0.01 | 0.1 | 0.1 |
| 3 | $10^{-4}$ | 50 | 50 | 500000 | 0.01 | 0.01 | 0.1 | 0.1 |
| | $10^{-2}$ | 50 | 500 | 50000 | 0.01 | 0.01 | 0.1 | 0.1 |
| | $10^{0}$ | 50 | 5000 | 5000 | 0.01 | 0.01 | 0.1 | 0.1 |
| 4 | $10^{-4}$ | 5 | 5 | 50000 | 0.01 | 0.01 | 0.05 | 0.05 |
| | $10^{-2}$ | 5 | 50 | 5000 | 0.01 | 0.01 | 0.05 | 0.05 |
| | $10^{0}$ | 5 | 500 | 500 | 0.01 | 0.01 | 0.05 | 0.05 |
| 5 | $10^{-4}$ | 5 | 5 | 50000 | 0.01 | 0.01 | 0.25 | 0.25 |
| | $10^{-2}$ | 5 | 50 | 5000 | 0.01 | 0.01 | 0.25 | 0.25 |
| | $10^{0}$ | 5 | 500 | 500 | 0.01 | 0.01 | 0.25 | 0.25 |
| 6 | $10^{-4}$ | 5 | 5 | 50000 | 0.05 | 0.05 | 0.1 | 0.1 |
| | $10^{-2}$ | 5 | 50 | 5000 | 0.05 | 0.05 | 0.1 | 0.1 |
| | $10^{0}$ | 5 | 500 | 500 | 0.05 | 0.05 | 0.1 | 0.1 |
| 7 | $10^{-4}$ | 5 | 5 | 50000 | 0.025 | 0.025 | 0.1 | 0.1 |
| | $10^{-2}$ | 5 | 50 | 5000 | 0.025 | 0.025 | 0.1 | 0.1 |
| | $10^{0}$ | 5 | 500 | 500 | 0.025 | 0.025 | 0.1 | 0.1 |

**Table S2: Measured stiffness of the BM** Mean Young's modulus at each measured position and mean Young's modulus over all positions, based on treatment group. $P$ values corresponding to the significance levels indicated in Fig. 5B. Pairwise t-test with Bonferroni correction for multiple testing.

| | | | | Young's modulus $E$ (kPa) | | adjusted $P$ value | |
|---|---|---|---|---|---|---|---|
| Animal | Condition | Position | $n$ | mean | sd | Position 1 | Position 2 |
| 1 | BBN | 1 | 20 | 46 | 27 | — | — |
| | | 2 | 37 | 100 | 47 | $7.42 \times 10^{-4}$ | — |
| | | 3 | 32 | 133 | 32 | $3.30 \times 10^{-10}$ | $3.10 \times 10^{-1}$ |
| 2 | BBN | 1 | 31 | 20 | 8 | — | — |
| | | 2 | 43 | 121 | 44 | $2.37 \times 10^{-20}$ | — |
| | | 3 | 39 | 125 | 50 | $4.46 \times 10^{-21}$ | 1 |
| 3 | BBN | 1 | 79 | 110 | 55 | — | — |
| | | 2 | 41 | 147 | 59 | $1.38 \times 10^{-3}$ | — |
| 4 | BBN | 2 | 32 | 27 | 18 | — | — |
| | | 3 | 39 | 77 | 30 | — | $1.94 \times 10^{-4}$ |
| 5 | CTRL | 1 | 44 | 136 | 47 | — | — |
| | | 2 | 42 | 139 | 39 | 1 | — |
| | | 3 | 40 | 104 | 37 | $1.38 \times 10^{-1}$ | $4.42 \times 10^{-2}$ |
| 6 | CTRL | 1 | 42 | 99 | 37 | — | — |
| | | 2 | 38 | 126 | 50 | $7.67 \times 10^{-1}$ | — |
| | | 3 | 46 | 120 | 40 | 1 | 1 |
| 7 | CTRL | 1 | 51 | 114 | 37 | — | — |
| | | 2 | 48 | 139 | 53 | $6.52 \times 10^{-1}$ | — |
| | | 3 | 46 | 101 | 31 | 1 | $2.67 \times 10^{-3}$ |
| 1–4 | BBN | all | 393 | 97 | 57 | | |
| 5–7 | CTRL | all | 397 | 120 | 44 | | |

**Table S3: Overview of animals used in this study** $n$: number of animals in total

| Experiment | weeks post BBN | $n$ | Group | | Comment |
|---|---|---|---|---|---|
| | | | BBN | CTRL | |
| Establishing treatment timeline | 2–11 | 16 | 16 | 0 | 4 mice each at 2, 5, 7 and 11 weeks |
| Histopathology 11 weeks | 11 | — | — | — | from 11 week sampling point timeline establishing |
| Histopathology 4 weeks | 4 | 12 | 6 | 6 | |
| SPIM imaging 11 weeks | 11 | 16 | 8 | 8 | |
| SPIM imaging 4 weeks | 4 | 36 | 18 | 18 | |
| AFM | 4 | 7 | 4 | 3 | |
| Total | 2–11 | 87 | 52 | 35 | |

### Supplementary Information

#### Elastic theory of wrinkling in multilayered tissues

In the computer simulations, we observed a transition between two morphologically distinct regimes of folding and wrinkling in the three-layer mucosa (Fig. 4D and S6A), depending on the relative stiffness of the epithelium (EP) and lamina propria (LP). If the passively deforming LP is much stiffer than the expanding EP ($E_{\mathrm{EP}} \ll E_{\mathrm{LP}}$) but softer than the basement membrane (BM), it is the BM alone that buckles and folds in between the EP and LP, which then act as compliant media for the BM, a regime we term "medium-like" deformation (Fig. 4D, upper left inset, and Fig. S6B). For a considerably stiffer EP ($E_{\mathrm{EP}} \gg E_{\mathrm{LP}}$), on the other hand, the EP deforms together with the BM like an expanding stiff plate on a compliant substrate, a regime we term "plate-like" deformation (Fig. 4D, lower right inset, and Fig. S6C). We propose that these two regimes and the transition between them provide a mechanical basis for structural changes in the mucosa that could be linked with the onset of bladder cancer formation through changes is tissue stiffness or thickness.

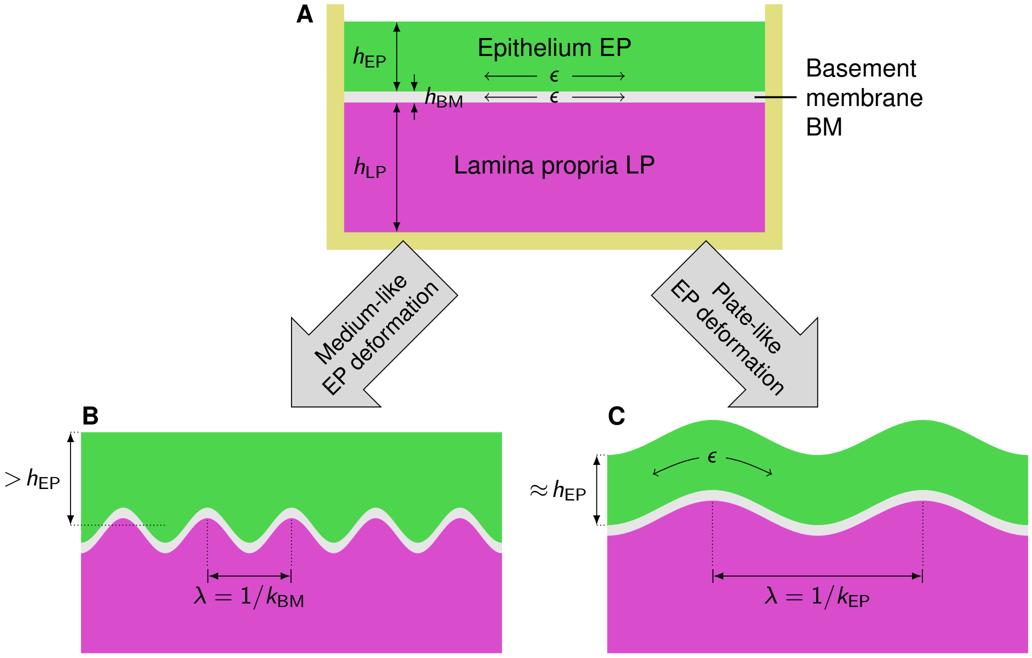


**Figure S6: Elastic theory of mucosa folding morphologies in 2D cross sections. A** Initial, flat, unstressed configuration of the mucosa, represented by a three-layer elastic continuum. The epithelium (EP) and the basement membrane (BM) are assumed to expand in plane, the lamina propria (LP) is not. **B** In "medium-like" deformation, the EP acts like an elastic medium similarly to the LP, sandwiching the thin BM, which folds between them upon expansion. The BM buckles and folds with wavelength $\lambda = 1/k_{\mathrm{BM}}$, as it cannot expand horizontally, straining the EP and LP vertically. If the EP overgrows in parallel to the BM and is also prevented from horizontal expansion, it thickens vertically. **C** In "plate-like" deformation, the EP buckles together with the BM like a composite plate on an elastic substrate, the LP. Since the wavelength is proportional to the plate thickness, it is larger, $\lambda \approx 1/k_{\mathrm{EP}}$. The undulating EP can attain its expanded length with lower horizontal compression, and therefore does not thicken substantially.

The existence of two regimes can be understood from the stability theory of elastic layered media, which goes back to the pioneering work of Biot [1]. Here, we derive the boundary separating them (Fig. 4D, black dashed line) analytically. Consider a 2D vertical cross section of the mucosa, and its small-deformation buckling behavior upon differential expansion. The EP, BM and LP layers are assumed to have unstrained initial thicknesses $h_{\mathrm{EP}}$, $h_{\mathrm{BM}}$ and $h_{\mathrm{LP}}$, and Young's moduli $E_{\mathrm{EP}}$, $E_{\mathrm{BM}}$ and $E_{\mathrm{LP}}$, with $h_{\mathrm{BM}} \ll h_{\mathrm{EP}}, h_{\mathrm{LP}}$ and $E_{\mathrm{BM}} \gg E_{\mathrm{EP}}, E_{\mathrm{LP}}$ (Fig. S6A). The EP and BM are assumed to expand horizontally by a volumetric growth strain $\epsilon > 0$, caused by proliferation of the epithelial cells and the membrane at the onset of cancer formation, for example. The 2D elastic energy per unit area of medium-like deformation is then given by the sum of four terms that account for vertical deformation of the EP and LP (which both act like an elastic half-space), bending of the BM, and compression of the volumetrically expanding EP. Assuming that all layers are incompressible with a Poisson ratio of 1/2, it reads [2]

$$U_{\mathrm{m}} = \frac{\epsilon}{3}\left(\frac{E_{\mathrm{EP}}}{\pi k_{\mathrm{BM}}} + \frac{4}{3}E_{\mathrm{BM}}h_{\mathrm{BM}}^3\pi^2 k_{\mathrm{BM}}^2 + \frac{E_{\mathrm{LP}}}{\pi k_{\mathrm{BM}}} + 2E_{\mathrm{EP}}h_{\mathrm{EP}}\epsilon\right) \tag{S1}$$

where

$$k_{\mathrm{BM}} = \frac{(3E_{\mathrm{LP}}/E_{\mathrm{BM}})^{1/3}}{2\pi h_{\mathrm{BM}}} \tag{S2}$$

is the wavenumber (inverse wavelength, $k = 1/\lambda$) of the undulating basement membrane after it has exceeded the critical planar compression. In plate-like deformation, however, the EP trades in compression for bending at longer wavelengths, because the latter becomes energetically favorable. The fourth energy term therefore is replaced by plate bending of the EP, which results in

$$U_{\mathrm{p}} = \frac{\epsilon}{3}\left(\frac{4}{3}E_{\mathrm{EP}}h_{\mathrm{EP}}^3\pi^2 k_{\mathrm{EP}}^2 + \frac{4}{3}E_{\mathrm{BM}}h_{\mathrm{BM}}^3\pi^2 k_{\mathrm{EP}}^2 + \frac{E_{\mathrm{LP}}}{\pi k_{\mathrm{EP}}}\right). \tag{S3}$$

Here,

$$k_{\mathrm{EP}} = \frac{(3E_{\mathrm{LP}}/E_{\mathrm{EP}})^{1/3}}{2\pi h_{\mathrm{EP}}} \tag{S4}$$

is the wavenumber that effectively results from EP bending on the LP substrate alone in good approximation, because $h_{\mathrm{BM}} \ll h_{\mathrm{EP}}$. The boundary of the two regimes can be found by equating the two elastic energy densities, $U_{\mathrm{m}} = U_{\mathrm{p}}$. For notational convenience, we now introduce the terms

$$r_{\mathrm{EP}} = \left(\frac{E_{\mathrm{EP}}}{E_{\mathrm{BM}}}\right)^{1/3}, \quad r_{\mathrm{LP}} = \left(\frac{E_{\mathrm{LP}}}{E_{\mathrm{BM}}}\right)^{1/3} \quad \text{and} \quad \widehat{h} = \frac{h_{\mathrm{EP}}}{h_{\mathrm{BM}}}. \tag{S5}$$

After some simplifications, the energy balance reads

$$3c\left(\widehat{h}\, r_{\mathrm{EP}} + \frac{1}{3\widehat{h}^2\, r_{\mathrm{EP}}^2} - 1\right) r_{\mathrm{LP}}^3 = 2\left(3\epsilon\widehat{h}\, r_{\mathrm{LP}} + c\right) r_{\mathrm{EP}}^3 \tag{S6}$$

where $c = 3^{2/3}$. Eq. S6 can be expressed in the form of a depressed cubic polynomial in $r_{\mathrm{LP}}$:

$$r_{\mathrm{LP}}^3 + p r_{\mathrm{LP}} + q = 0 \tag{S7}$$

with coefficients

$$\begin{aligned} p &= -\frac{2\epsilon\widehat{h}d}{c} < 0,\\ q &= -\frac{2d}{3} < 0,\\ d &= \frac{r_{\mathrm{EP}}^3}{\widehat{h}\, r_{\mathrm{EP}} + 1/(3\widehat{h}^2\, r_{\mathrm{EP}}^2) - 1} > 0. \end{aligned} \tag{S8}$$

Cardano's formula then yields the morphological phase boundary, as shown in Figs. 4D and S7, in explicit form:

$$\frac{E_{\mathrm{LP}}}{E_{\mathrm{BM}}} = r_{\mathrm{LP}}^3 = \left(u_1^{1/3} + u_2^{1/3}\right)^3 \quad \text{with} \quad u_{1,2} = -\frac{q}{2} \pm \sqrt{\left(\frac{q}{2}\right)^2 + \left(\frac{p}{3}\right)^3}. \tag{S9}$$

In Fig. S7A, we plot the morphological boundary in the plane of stiffness ratios between the three layers for different amounts of volumetric expansion, $\epsilon$. For larger $\epsilon$, the boundary shifts toward stiffer lamina

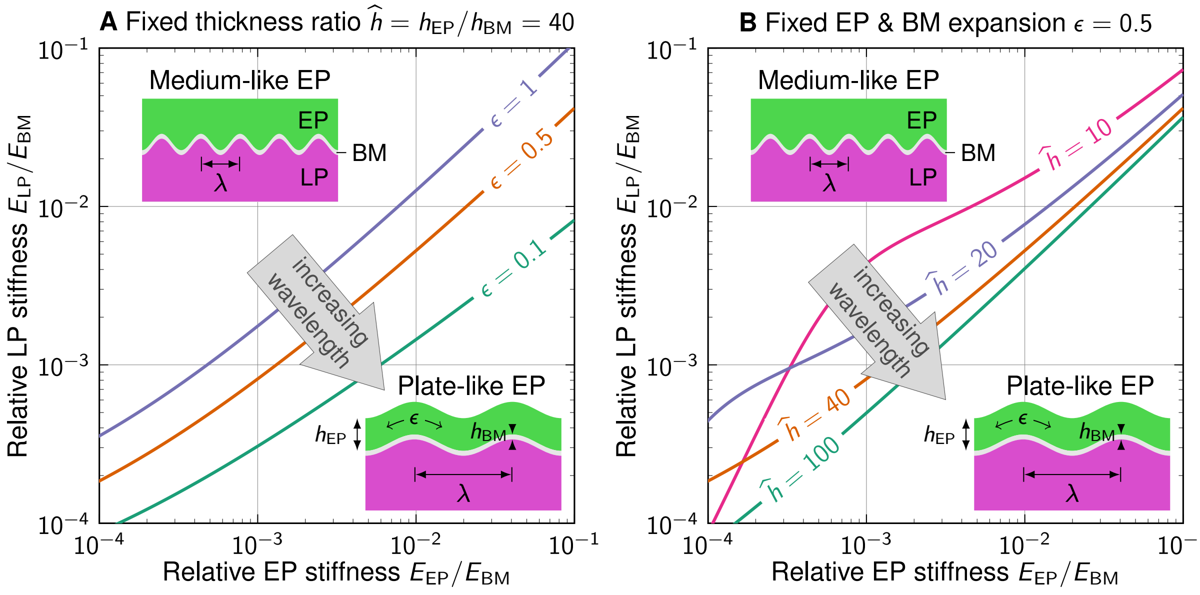


**Figure S7: Morphological transition between different epithelium folding regimes. A** Theoretical boundaries between medium-like and plate-like deformations of the expanding epithelium at a fixed ratio of layer thicknesses, for different volumetric expansions. **B** Theoretical boundaries at a fixed volumetric expansion, for different thickness ratios.

propria, making the plate-like bending of the epithelium somewhat more favorable at large overgrowth of the upper two layers. Yet, both morphologies coexist also in the small-strain limit, in which the boundary lies at $E_{LP}/E_{BM} \to -q$ for $\epsilon \to 0$, which can be written as

$$\frac{E_{LP}}{E_{EP}} \to \frac{2}{3(f-1)+f^{-2}} \quad \text{for} \quad \epsilon \to 0 \quad \text{with} \quad f = \frac{h_{EP}}{h_{BM}} \left(\frac{E_{EP}}{E_{BM}}\right)^{1/3}. \tag{S10}$$

In Fig. S7B, the morphological regime boundary is plotted at different thickness ratios. Thicker epithelia (relative to the BM) allow for a larger region in the stiffness plane in which the medium-like deformation is favorable. Taking the limit of a very thick EP or thin BM for the general solution (Eq. S9) yields a linear regime boundary at

$$\frac{E_{LP}}{E_{EP}} \to \frac{2}{3}\sqrt{2\epsilon^3} \quad \text{for} \quad \hat{h} \to \infty. \tag{S11}$$

Note that, in all these results, the layer stiffnesses and thicknesses occur only in the form of ratios, not absolute numbers, which implies that this mechanical transition can be found across the scales. This may provide a common mechanical basis for the explanation of cancer formation in different multilayer tissues and across species. Medium-like deformation crumples the basement membrane at short wavelengths, potentially pushing it beyond the yield point at which plastic structural damage is dealt to the cellular structure or ECM. These short-wavelength folds in the basement membrane, together with irreversible damage and plastic stress localization, may be a mechanical explanation for the onset of carcinomas in situ.
